# Greater hypoxic burden predicts weaker coordination between brain pulsation and CSF flow on 7T MRI independent of non-hypoxic arousals: Implications for glymphatic activity

**DOI:** 10.64898/2026.04.25.720853

**Authors:** Gawon Cho, Korey Kam, Anne Chen, Yutong Mao, Helene Benveniste, Adam Mecca, Daphne I. Valencia, Katarina R. Martillo, Sarah S. Chu, Ankita Kumar, Omonigho M. Bubu, Ricardo S. Osorio, Indu Ayappa, Xiao Liu, Brienne Miner, Andrew W. Varga

## Abstract

**Study Objectives:** Obstructive sleep apnea (OSA) is a risk factor for neurodegeneration, and glymphatic impairment may be one mechanistic pathway. Anti-phase coordination between brain pulsations and CSF flow, reflecting compensatory CSF displacement during each vascular pulsation cycle, supports glymphatic activity. This study examined whether hypoxic burden and sleep fragmentation, two distinct OSA pathologies, are differentially associated with brain pulsation-CSF flow dynamics.

**Methods:** This cross-sectional study included 28 individuals with newly identified OSA and 8 without OSA. Participants completed in-lab polysomnography or WatchPAT testing, providing measures of hypoxic burden, quantified as time below 90% oxygen saturation (T90), and non-hypoxic sleep fragmentation, quantified as respiratory effort-related arousals (RERAs). Participants also completed 7T resting-state functional MRI to estimate global BOLD-CSF, defined as anti-phase cross-correlation between blood-oxygen-level-dependent (BOLD) signal and CSF inflow. Associations were examined using correlation and hierarchical regression. Exploratory analyses examined region-specific BOLD-CSF and brain pulsation strength, quantified as BOLD amplitude.

**Results:** Greater T90 was associated with weaker global BOLD-CSF, independent of RERAs and covariates (β=0.08,p=0.03). Greater T90 was also associated with higher BOLD amplitude across temporal, frontal, and parietal regions, but this elevation in amplitude was not accompanied by stronger region-specific BOLD-CSF coupling (β=0.001,p>0.05). In contrast, among regions where BOLD amplitude was not associated with T90, greater BOLD amplitude predicted stronger region-specific BOLD-CSF (β=−0.004,p<0.001). RERAs were not associated with global BOLD-CSF or BOLD amplitude.

**Conclusions:** In OSA, hypoxic burden may be the primary feature associated with impaired brain pulsation and CSF dynamics that support glymphatic activity. These alterations may be pronounced in the temporal lobe, where elevated pulsations were uncoupled from compensatory CSF displacement.

## 1. Introduction

Obstructive sleep apnea (OSA) is a disorder characterized by recurrent upper airway obstruction, and affects nearly one billion adults worldwide.^1^ OSA is a robust risk factor for multiple neurodegenerative conditions including Alzheimer’s disease and related dementias (ADRD),^2^ with glymphatic impairment representing a potential mechanistic pathway.^3-5^ The glymphatic system is a recently proposed brain waste clearance pathway, which is thought to transport metabolic waste products out of the brain via bulk cerebrospinal fluid (CSF) flow through perivascular spaces.^6-11^ A key driver of glymphatic transport is anti-phase coordination between brain pulsations (defined as rhythmic expansions and contractions of the vasculature and surrounding tissue) and CSF flow.^12-14^ Because the brain must maintain constant intracranial volume, blood inflow and displacement during each pulsation cycle are accompanied by reciprocal CSF volume change, generating the anti-phase fluid motion necessary for solute transport.^12-15^ Prior research has shown that individuals with OSA exhibit weaker coordination between brain pulsations and CSF flow, with greater discoordination observed at higher levels of apnea-hypopnea index (AHI).^16-18^ However, AHI reflects a composite of two distinct consequences of intermittent upper airway closure in OSA, namely intermittent hypoxemia and respiratory effort-related sleep fragmentation.^1^ Thus, the respective contributions of hypoxic burden and respiratory effort-related sleep fragmentation to the coordination between brain pulsations and CSF flow, independent of one another, remain unknown.

In addition to the coordination between brain pulsations and CSF flow, the strength of brain pulsations themselves is also central to glymphatic transport.^13,15^ Increases in the amplitude of rhythmic expansions and contractions of the vasculature and surrounding tissue can produce larger perivascular space volume changes, which can translate into stronger pulsatile CSF flow.^14^ Prior animal research has shown vascular pulsation amplitude to be positively associated with perivascular solute transport.^14^ Nonetheless, whether the two distinct pathological mechanisms of OSA, namely hypoxic burden and respiratory effort-related sleep fragmentation, are differentially associated with brain pulsation strength remains unclear.

Distinguishing between potential impact of hypoxic burden and respiratory effort-related sleep fragmentation is important because the two OSA features may differentially modulate the glymphatic system. Hypoxic burden, one variable for which is time spent below 90% oxygen saturation,^19^ may induce hypoxic injury to the vasculature, leading to vascular stiffening,^20^ thereby interfering with brain pulsations and their coordination with CSF flow needed for improved glymphatic transport. Regions with high metabolic demand, such as the frontotemporal cortex, may be especially vulnerable to hypoxic injury of cerebral vessels,^21^ which could translate into greater alterations in brain pulsations and their coordination with CSF flow. On the other hand, respiratory effort-related sleep fragmentation without hypoxemia can be captured by respiratory effort-related arousals (RERAs), defined as respiratory events that lead to arousal without a significant drop in peripheral oxygen saturation.^22,23^ Such incidences of sleep fragmentation may disrupt oscillations in norepinephrine tone, which contribute to rhythmic vasoconstriction and vasodilation of the cerebrovasculature that drive glymphatic activity.^11,14,24^

To address these gaps, this study examined the associations of hypoxic burden and non-hypoxic sleep fragmentation with the coordination between brain pulsations and CSF flow, with particular emphasis on distinguishing their independent associations. In addition, the study examined the associations of hypoxic burden and non-hypoxic sleep fragmentation with region-specific brain pulsation strength, quantified as pulsation amplitude. From a clinical perspective, these findings may help clarify whether interventions that preferentially reduce hypoxemia, improve sleep continuity, or target both processes are more likely to improve fluid dynamics that support glymphatic activity.

## 2. Methods

### 2.1. Sample

This cross-sectional study examined 36 individuals aged 55 to 76 years. The sample included 22 adults with a new clinical diagnosis of OSA and not yet receiving treatments recruited from the Mount Sinai Integrative Sleep Center and 14 community-dwelling, cognitively normal older adults who completed in-lab polysomnography at the Mount Sinai School of Medicine. While the community-recruited subjects did not report any sleep complaints, OSA (AHI4% > 5/hour) was incidentally identified in 6 out of the 14 subjects, resulting in a final sample of 28 individuals with OSA and 8 OSA-free controls. Exclusion criteria included having any self-identified cognitive complaints, sleep disorders other than OSA, sleep duration of less than 5 hours or greater than 10 hours per night based on actigraphy, significant neurological or psychiatric disorders, chronic usage of any psychoactive drugs for a duration over 8 weeks aside from single selective serotonin reuptake inhibitors, any unstable comorbid conditions, and metal in the body. The final sample included 36 participants who met the study’s eligibility criteria (please see Supplementary Table 1 for handling of missing data and participant flow). The current study protocols were approved by the Mount Sinai Institutional Review Board, and additional analyses of de-identified data were classified as non-human subjects research by the Yale Institutional Review Board.

**Table 1.** Sample characteristics.

| Characteristics | N (%) or Median (IQR) |  |  |
| --- | --- | --- | --- |
|  | Full sample | OSA (N=28) | Controls (N=8) |
| Age | 66 (62-71) | 65.5 (61-71) | 67.5 (65-70) |
| Female sex | 21 (58.3%) | 14 (50) | 7 (87.5) |
| White | 24 (66.7%) | 17 (60.7) | 7 (87.5) |
| BMI | 30.0 (26.9-33.5) | 30.5 (28.2-35.6) | 24.2 (22.0-29.8) * |
| Epworth Sleepiness Scale | 5 (3-8) | 5 (3-8.5) | 4 (2.5-6) |
| AHI3% (per hour) | 29.9 (9.3-49.0) | 33.6 (23.0-54.7) | 2.9 (1.9-4.2) * |
| AHI4% (per hour) | 22.6 (6.3-38.7) | 26.6 (17.7-47.4) | 1.7 (1.2-3.5) * |
| T90 (in minutes) | 5.7 (1.1-21.5) | 7.5 (3.0-37.1) | 0.8 (0.3-1.2) * |
| Non-hypoxic RERAs<sup>1</sup> (per hour) | 1.3 (0.5-2.9) | 1.7 (0.7-2.9) | 1.1 (0.5-3.4) |
| Total sleep time (in minutes) | 387.5 (346.8-425) | 394.8 (361.5-431.5) | 348.3 (334.5-378.1) |
| Hypertension | 15 (41.7) | 12 (42.9) | 3 (37.5) |
| MoCA | 27 (25-29) | 27 (25-28.5) | 29 (27-30) |

<sup>1</sup> RDI – AHI4%
\* Denotes statistically different group differences
*Acronyms.* AHI4%=apnea-hypopnea index using the 4% criteria; BMI=body mass index; IQR=Interquartile range; MoCA=Montreal Cognitive Assessment; RERA=respiratory effort-related arousal index; SD=standard deviation; T90=time of sleep spent under 90% oxygen saturation

### 2.2. Diagnostic sleep studies

Participants completed either in-laboratory polysomnography or WatchPAT-based home sleep test, a clinically validated method for assessing OSA that has shown excellent agreement with in-laboratory polysomnography.^25-27^ The procedures for each modality are described below.

### 2.2.a. Polysomnography

Fourteen participants recruited from the community completed an overnight, attended, in-laboratory polysomnography at the Mount Sinai Integrative Sleep Center using 2.2.a.

Compumedics E-series (Melbourne, Australia).^28^ Participants were connected to polysomnography and the lights off time was scheduled to approximate their self-reported habitual bedtime. Signal acquisition included electroencephalography, electro-oculography, electromyography, a nasal cannula, pressure transducer, oral thermistor, rib or abdomen impedance plethysmography, single-channel electrocardiography, and pulse oximetry. All polysomnography recordings were scored by 1 or 2 registered polysomnographic technologists according to the American Academy of Sleep Medicine (AASM) standard criteria,^28^ and one sleep technologist (I.A.) reviewed all sleep studies for the consistency of scoring.

For all participants, hypoxic burden was defined as time spent below 90% oxygen saturation, henceforth referred to as T90. T90 was calculated as the cumulative duration, in minutes, during which peripheral oxygen saturation measured by pulse oximetry remained below 90%. In addition, non-hypoxic sleep fragmentation was defined using respiratory effort-related arousals (RERAs), a tally of arousals linked to respiratory events per hour without significant peripheral oxygen desaturation. RERAs were derived by subtracting the apnea-hypopnea index using the 3% desaturation criterion (AHI3%) from AHI3%a, defined as the apnea-hypopnea index based on events associated with either ≥3% oxygen desaturation or arousal.^29,30^ Here, apneas were defined as events with absence of airflow for ≥10 seconds. Hypopneas were defined as events with a reduction in breathing amplitude by ≥30% for ≥10 seconds, accompanied by a ≥3% decline in blood oxygen saturation, regardless of arousal status. Hypopnea 3A was defined as an event with a reduction in breathing amplitude by ≥30% for ≥10 seconds, terminating in either a ≥3% decline in blood oxygen saturation or an arousal. AHI3% was defined as the sum of apneas and hypopneas with ≥3% desaturation divided by total sleep time in hours. AHI3%a was defined as the sum of apneas and hypopneas with ≥3% desaturation or arousal divided by total sleep time in hours.^29,30^ While a ≥4% decline in blood oxygen saturation is commonly used to identify hypopneas, we used the 3% desaturation criterion to more clearly distinguish non-hypoxic RERAs from desaturation-associated respiratory events.

### 2.2.b. WatchPAT Home Sleep Test

Twenty-two participants newly diagnosed with OSA who were not yet receiving treatment were recruited from the Mount Sinai Sleep Center and wore WatchPAT ONE devices (Zoll, Germany).^19^ WatchPATs measure sleep duration based on wrist actigraphy, snoring and chest breathing excursions using a wireless thoracic sensor with a microphone, and pulse oximetry and peripheral arterial tonometry (PAT) based on a finger probe.^19^ Changes in PAT and movement were used to capture increases in sympathetic tone that are linked to respiratory effort-related arousals.^19^ Automated respiratory event scoring algorithms were used to derive PAT-based measures of T90, respiratory disturbance index (henceforth referred to as PAT-derived RDI [pRDI], which includes the sum of total apneas, hypopneas, and RERAs), and AHI3%. Non-hypoxic RERAs for participants who wore WatchPATs were calculated by subtracting AHI3% from pRDI, following the AASM manual and prior research.^22,23^ Prior research shows excellent agreement in sleep and respiratory event metrics between in-lab polysomnography and WatchPAT for T90 (intraclass correlation coefficient [ICC]=0.86),^19,25-27^ WatchPAT-derived pRDI showed excellent agreement with AHI3%a from in-lab polysomnography (ICC=0.76; Supplementary Table 1). Likewise, WatchPAT-derived AHI3% showed excellent agreement with AHI3% from in-lab polysomnography (ICC=0.76; Supplementary Table 1).

### 2.3. 7T MRI Protocol

Participants completed 9 to 15 minutes of resting-state functional MRI on a 7T scanner (Magnetom, Siemens, Erlangen, Germany) with a 32-channel head coil (repetition time [TR]=1.5 seconds, echo time [TE]=0.025, flip angle [FA]=55°, iPAT acceleration factor=3, bandwidth=33.921/pixel, 1.5 mm isotropic resolution). For the blood oxygen level dependent signal (BOLD) sequence, the bottom slice was aligned to the 4^th^ ventricle to capture CSF inflow, consistent with prior research.^31^ In addition, whole-brain T1-weighted gradient echo (MP2RAGE) anatomical images were acquired (TR=4500 ms, TE=3.37 ms, TI1=1000s and 3200 ms, FA=4 and 5°, iPAT acceleration factor=3, bandwidth =130 Hz/pixel, 0.7 mm isotropic resolution). Processing steps for resting-state scans included deobliquing, motion-correcting, skull-stripping, spatial smoothing, grand mean-scaling, detrending of linear and quadratic trends, and registration to T1.

### 2.3.a. Coordination between brain pulsation and CSF flow at resting-state

To quantify the coordination between *global* brain pulsations and CSF inflow, we calculated anti-cross-correlation strength between the global blood oxygen level dependent signal captured in the gray matter and CSF inflow captured in the 4th ventricle^12^ (henceforth referred to as gBOLD-CSF). Anti-phase cross-correlation was used to quantify coordination, based on the premise that blood inflow and displacement during each pulsation cycle must be accompanied by reciprocal CSF movement into or out of the cranium, producing tight temporal coupling between brain pulsations and CSF flow in healthy brains.^12^ Thus, more negative anti-cross-correlation values indicate putatively healthier anti-phase coordination between brain pulsations and CSF flow.^32-34^ Using processed resting-state scans in participants’ native space, global BOLD time series were derived as the average signal across all gray matter voxels using a gray matter mask, and CSF inflow time series were derived as the average signal across manually identified CSF voxels in the bottom slice aligned to the 4th ventricle.^35-37^ Both signals were filtered to the vasomotion frequency range, defined as <0.1 Hz, because brain pulsations in this range reflect slow oscillations in vascular tone driven by spontaneous smooth muscle activity in the microvasculature, which often show the greatest amplitude and have been found to exert the strongest effect on CSF transport,^13,15,31,38^ independent of heartbeat-generated pulsations at approximately 1 Hz. Each participant’s cross-correlation value at the 3-second lag, where the greatest sample-wide average anti-cross-correlation was identified, was designated as that participant’s gBOLD-CSF coupling strength.

In addition, we calculated a region-specific analogue of gBOLD-CSF, henceforth referred to as rBOLD-CSF, to quantify coordination between regional brain pulsations and CSF inflow. rBOLD-CSF was defined as the peak anti-cross-correlation between region-specific gray matter blood oxygen level dependent signals and CSF inflow captured in the 4th ventricle.^12^ This regional measure extends the same physiological framework used for gBOLD-CSF, in which blood inflow and displacement during each pulsation cycle must be accompanied by reciprocal CSF movement into or out of the cranium.^12^ Accordingly, more negative anti-cross-correlation values were interpreted as stronger and putatively healthier region-specific anti-phase coordination between brain pulsations specific to a region and CSF inflow.^32-34^ Region-specific gray matter BOLD time series were derived as the average within each Desikan-Killiany region.^39^ CSF inflow time series were calculated using the same approach as gBOLD-CSF. Subsequently, the anti-phase cross-correlation between each regional time series and the CSF signal at the 3-second lag, corresponding to the lag at which the global signal showed peak anti-phase cross-correlation, was used as that participant’s region-specific BOLD-CSF coupling strength.

### 2.3.b. Brain pulsation strength at resting-state

The strength of global brain pulsations was defined as the amplitude of gray matter-wide BOLD signals, henceforth referred to as gBOLD-amp. Following procedures used in previous studies, the 3dRSFC function in AFNI was applied to processed resting-state scans with a brain mask to generate voxelwise maps of resting-state fluctuation amplitude in the vasomotion frequency range, defined as <0.1 Hz.^13,15,16,31,38^ Here, resting-state fluctuation amplitude reflects the standard deviation of the voxelwise time series within the specified frequency range. gBOLD-amp was calculated by averaging the voxelwise amplitude values across all gray matter voxels. In addition, we calculated a regional analogue of gBOLD-amp, henceforth referred to as region-specific BOLD amplitude, or rBOLD-amp. rBOLD-amp was calculated by averaging voxelwise amplitude values within each Desikan-Killiany region.^39^

### 2.4. Statistical analysis

The primary analysis examined associations of gBOLD-CSF with T90 and non-hypoxic RERAs, respectively, to quantify the variance in coordination between global brain pulsations and CSF flow accounted for by each distinct type of OSA burden. Associations were examined using Pearson correlations and hierarchical linear regression models in which gBOLD-CSF coupling was modeled as a function of age, sex (female vs. male), sleep device (WatchPAT vs. in-lab polysomnogram), hypertension status (yes vs. no), non-hypoxic RERAs, and T90. The statistical significance of non-hypoxic RERAs and T90, as well as the incremental variance explained by inclusion of each term, was evaluated using the change in R^2^. Because RERAs include non-hypoxic arousals only, we examined the sensitivity of results by replacing RERAs with AHI3%a, which also includes hypoxic arousals, in Pearson correlation and hierarchical regression analyses.

Furthermore, to investigate whether OSA is associated with brain pulsation strength, we repeated the Pearson correlation and hierarchical regression analyses using gBOLD-amp as the outcome instead of gBOLD-CSF coupling. These models examined associations of gBOLD-amp with T90 and non-hypoxic RERAs, as well as the sensitivity of results to replacing RERAs with AHI3%a. Subsequently, to identify brain regions with increased susceptibility to OSA, we examined whether region-specific brain pulsation strength, measured as rBOLD-amp, was associated with T90 and non-hypoxic RERAs in each cortical region. Brain regions were then classified dichotomously according to whether rBOLD-amp was associated with T90, defining “T90-susceptible regions.” The same approach was used to define “RERA-susceptible regions.” Finally, we explored variations in the association between region-specific brain pulsation strength and region-specific coordination of brain pulsations with CSF flow between T90-susceptible and non-T90-susceptible regions. A mixed-effects model was used in which rBOLD-CSF was regressed on rBOLD-amp, dichotomous T90-susceptible region status, and their interaction, with brain region-level observations nested within individuals. An analogous mixed effects model was used to explore RERA-susceptible regions. All skewed variables identified using the Shapiro-Wilks test, which included T90, RERAs, AHI3%a, gBOLD-amp, rBOLD-amp, and rBOLD-CSF, were log-transformed to satisfy the condition of normality. One-sided p-values of Pearson correlation coefficients were derived from 10,000 permutation shuffles. Statistical significance was defined using a p-value of <0.05. All analyses were conducted in MATLAB R2024a and Stata/MP 17.0.

## 3. Results

Sample characteristics can be found in Table 1. Median age was 66 years [interquartile range,IQR=62-71], 58.3% of participants were female, and 66.7% were White. The median Montreal Cognitive Assessment score was 27 [IQR=25-29], and mean body mass index was 30.3 [5.5]. Participants had a median AHI4% of 22.6 events per hour [IQR = 6.3 to 38.7], a median RERA index of 1.3 events per hour [IQR = 0.5 to 2.9], and a median T90 of 5.7 minutes [IQR = 1.1 to 21.5]. Median of sleep duration was 387.5 minutes [IQR=346.8-425]. While the majority of characteristics were comparable between groups with and without OSA, the OSA group had fewer females and higher BMI, AHI3%, AHI4%, and T90.

Pearson correlation analysis showed that greater T90 was associated with weaker gBOLD-CSF coupling (Figure 1A; Pearson r=0.32, permutation p=0.029), as higher gBOLD-CSF values indicate weaker temporal coupling between gray matter pulsations and CSF flow. This association remained significant after adjustment for age, sex, sleep device, hypertension, and RERAs (Table 2; β=0.003, p=0.01). Hierarchical modeling further showed that adding T90 significantly improved the proportion of variance in gBOLD-CSF explained by the model (Table 2; ΔR^2^=0.18, p=0.01). Notably, neither non-hypoxic RERA’s nor the AHI3a were correlated with gBOLD CSF (RERAs: Figure 1B, Pearson r=0.02, p=0.46; AHI3a:Figure 1C, Pearson r=0.24, p=0.09) or explained significant variance in gBOLD-CSF beyond the base model of age, sex, sleep device, and hypertension (Table 2).

**Table 2.**
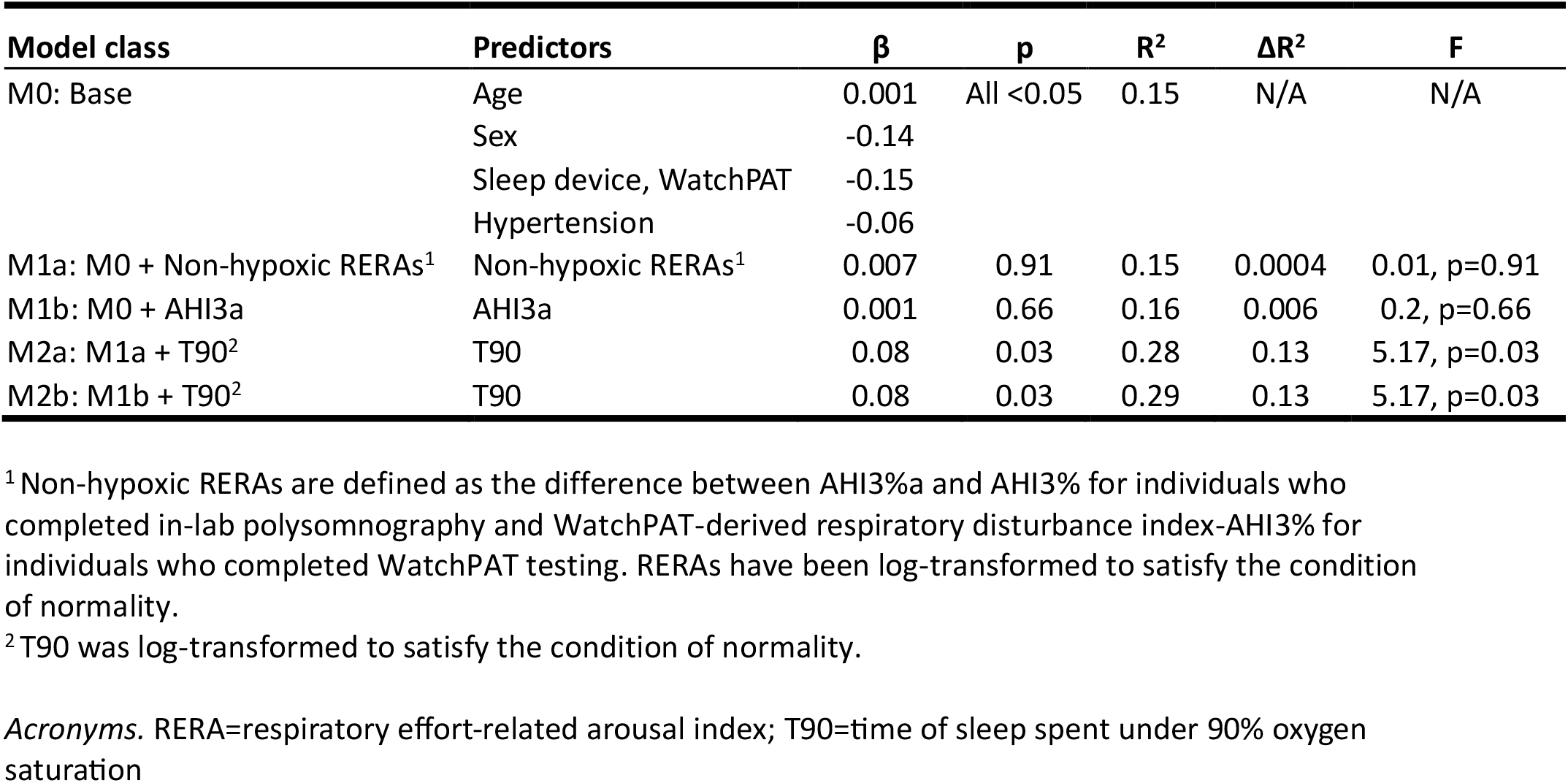
Association of T90 with gBOLD-CSF derived from a nested, hierarchical modeling approach.

**Figure 1.**
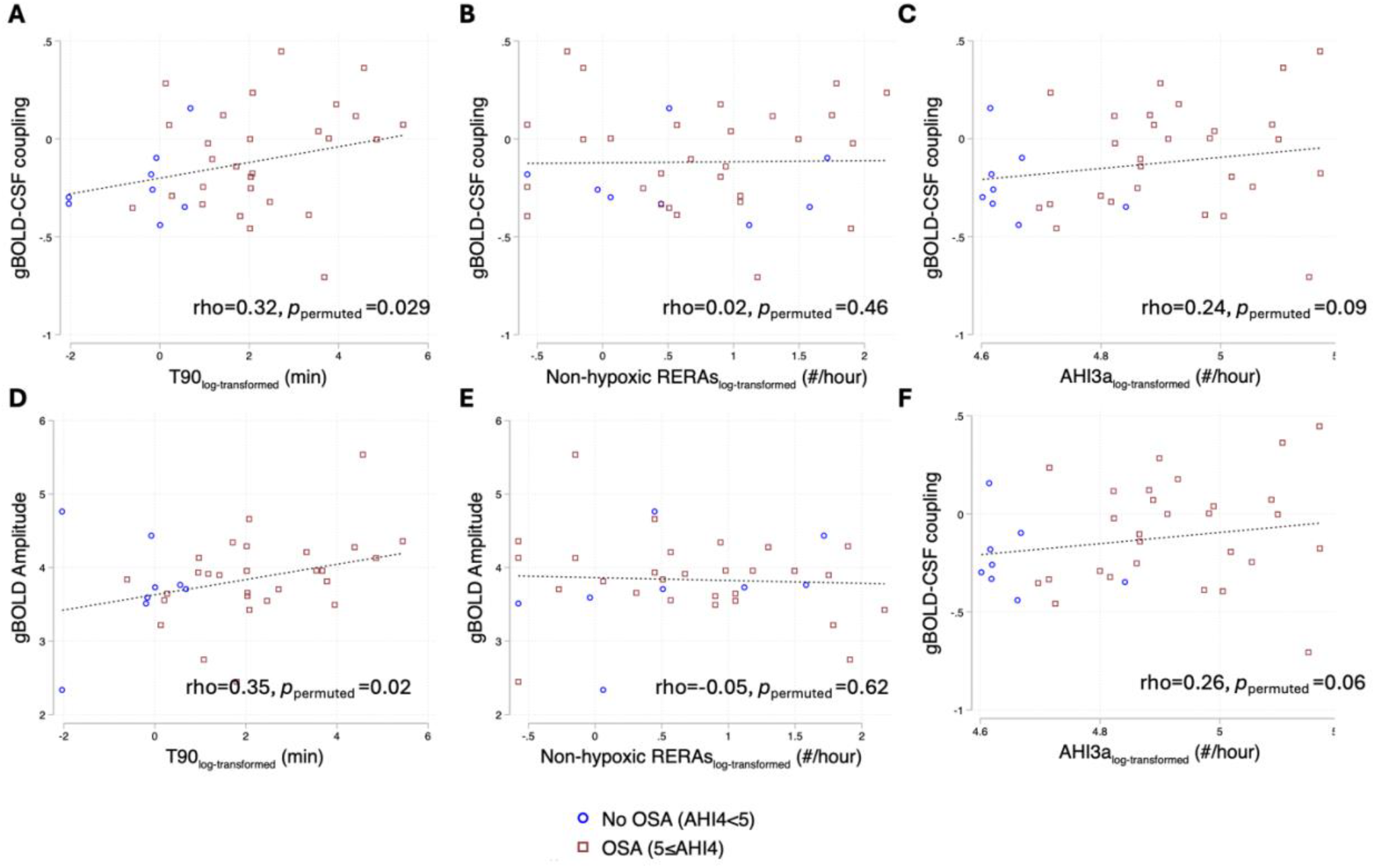
Pearson correlation of OSA characteristics with gBOLD-CSF coupling and gBOLD-amp *Acronyms*. AHI3a=apnea-hypopnea index using the 3% criteria + arousals; AHI4=apnea-hypopnea index using the 4% criteria; CSF=cerebrospinal fluid; gBOLD-amp=global gray matter pulsation strength; gBOLD-CSF coupling=coupling strength of global gray matter blood oxygen level dependent signal to CSF inflow; RERAs=respiratory effort-related arousals; T90=time of sleep spent under 90% oxygen saturation

Moreover, greater T90 was correlated with stronger global brain pulsation, quantified using gBOLD-amp (Figure 1D; Pearson r=0.35, permutation p=0.02), whereas non-hypoxic RERAs and AHI3%a were not (Figures 1E-F). The association between gBOLD-amp and T90 was attenuated and borderline significant after adjustments for covariates and RERAs (Table 3; β=0.12, p=0.06), and no longer significant in the sensitivity analysis involving AHI3%a instead of RERAs. Because the observed direction of the association was opposite to what we expected, we conducted further exploratory analyses to examine regional variations in the relationship between T90 and region-specific pulsation strength, measured as rBOLD-amp. Here, higher T90 was correlated with greater rBOLD-amp across the temporal lobe and multiple frontal and parietal regions, but not in the occipital cortex (Figure 2; region-specific estimates are provided in Supplementary Table 2). Within regions showing T90-associated elevations in rBOLD-amp, classified as “T90-susceptible regions”, rBOLD-amp was not associated with region-specific BOLD-CSF coupling strength (Colored in red in Figure 3; β=0.001, p>0.05), suggesting that elevated brain pulsation strength did not translate into stronger coordination with CSF flow. In contrast, among the “non-T90-susceptible regions”, where rBOLD-amp was not associated with T90, greater rBOLD-amp robustly predicted stronger region-specific BOLD-CSF coupling (Colored in black in Figure 3; β=−0.004, p<0.001; difference in β with “T90-susceptible regions”=0.005, p<0.001). RERAs were not associated with rBOLD-amp in any cortical regions (Supplementary Table 3).

**Table 3.** Association of T90 with gBOLD-amp derived from a nested, hierarchical modeling approach.

| Model class | Predictors | $\beta$ | p | R<sup>2</sup> | $\Delta R^2$ | F |
| --- | --- | --- | --- | --- | --- | --- |
| M0: Base | Age | -0.005 | All <0.05 | 0.09 | N/A | N/A |
|  | Sex | 0.12 |  |  |  |  |
|  | Sleep device, WatchPAT | -0.18 |  |  |  |  |
|  | Hypertension | -0.30 |  |  |  |  |
| M1a: M0 + Non-hypoxic RERAs<sup>1</sup> | Non-hypoxic RERAs<sup>1</sup> | -0.12 | 0.87 | 0.09 | 0.0008 | 0.03, p=0.87 |
| M1b: M0 + AHI3a | AHI3a | 0.009 | 0.23 | 0.13 | 0.04 | 1.52, p=0.23 |
| M2a: M1a + T90<sup>2</sup> | T90 | 0.17 | 0.05 | 0.20 | 0.12 | 4.18, p=0.0501 |
| M2b: M1b + T90<sup>2</sup> | T90 | 0.15 | 0.11 | 0.20 | 0.07 | 2.73, p=0.11 |

<sup>1</sup> Non-hypoxic RERAs are defined as the difference between AHI3%a and AHI3% for individuals who completed in-lab polysomnography and WatchPAT-derived respiratory disturbance index-AHI3% for individuals who completed WatchPAT testing. RERAs have been log-transformed to satisfy the condition of normality.
<sup>2</sup> T90 was log-transformed to satisfy the condition of normality.
*Acronyms.* RERA=respiratory effort-related arousal index; T90=time of sleep spent under 90% oxygen saturation

**Figure 2.**
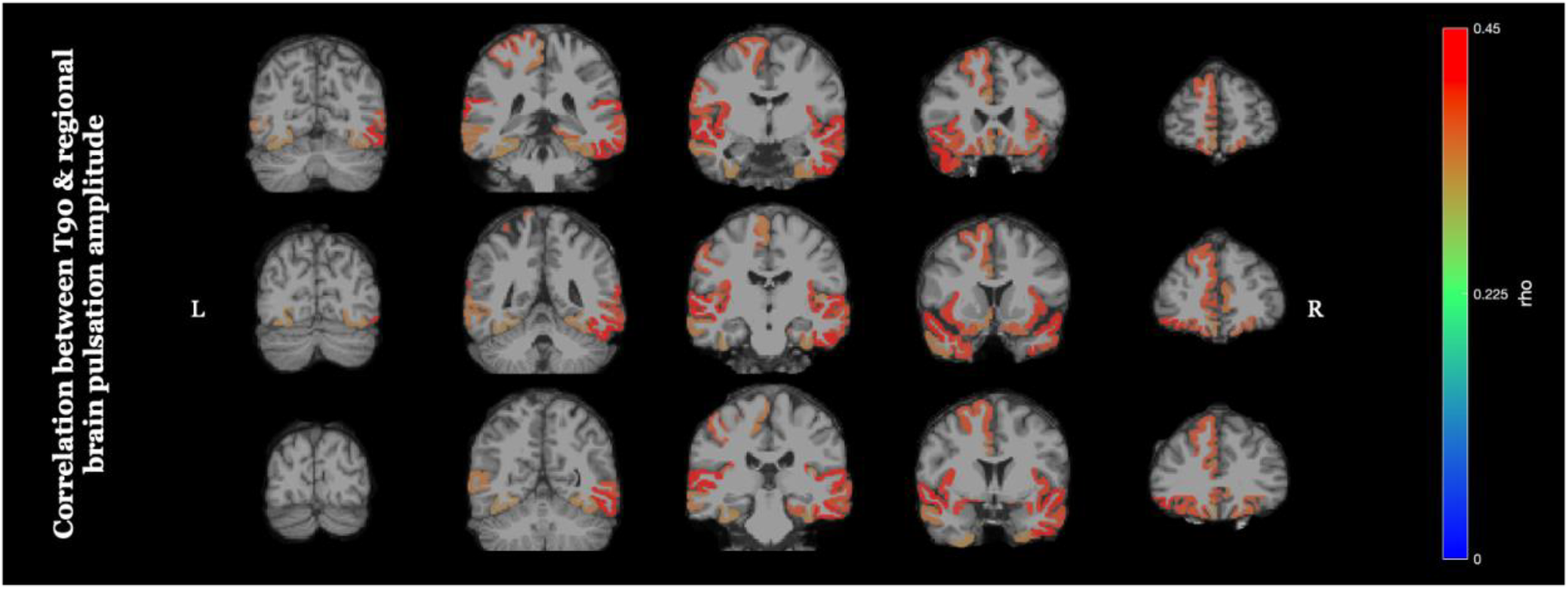
Association between T90 and coupling strength of regional gray matter pulsation amplitude T90 has been log-transformed to satisfy the condition of normality. Acronyms. r=Pearson correlation coefficient; T90=time spent under 90% oxygen saturation

**Figure 3.**
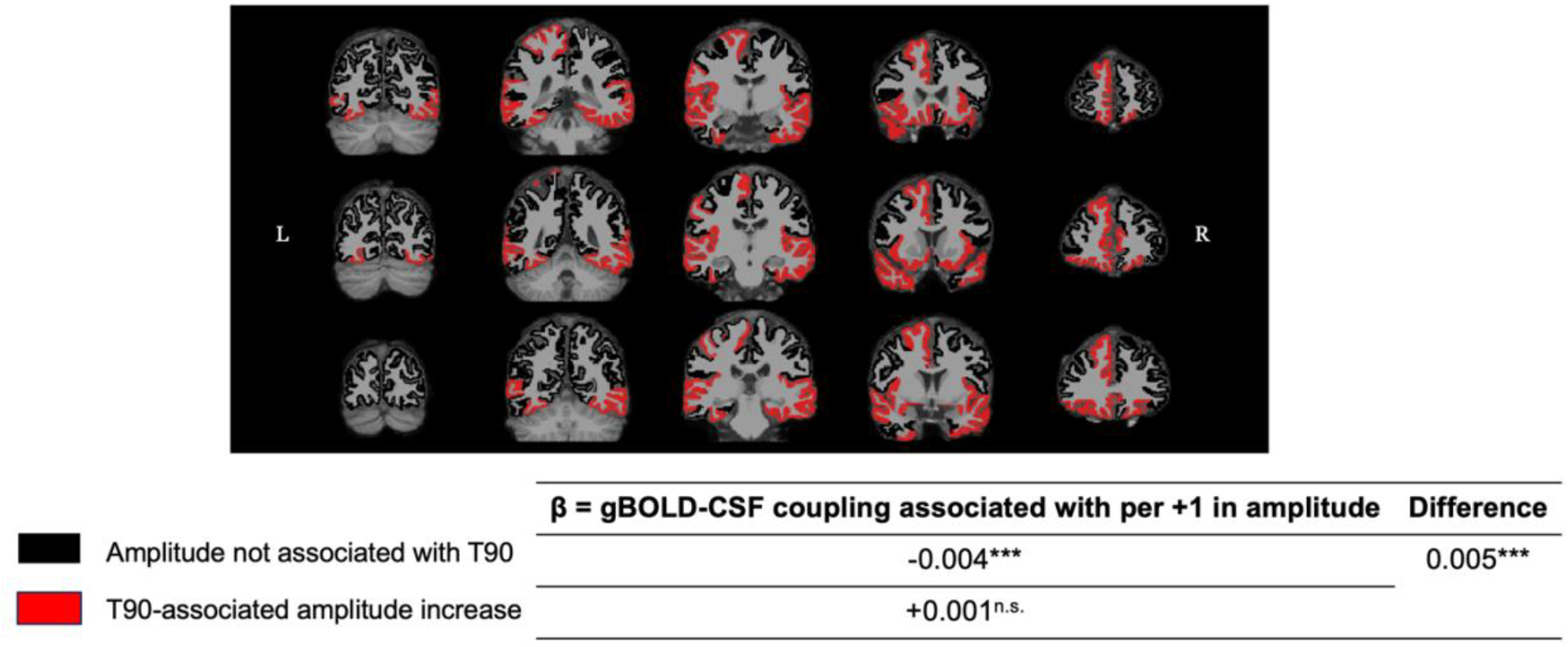
Association between amplitude and gBOLD-coupling by region Regional gBOLD-CSF coupling, regional gBOLD amplitude, and T90 have been log-transformed to satisfy the condition of normality. *** denotes a p-value of <0.001 The only way to say something about region-specific pulsations and their relationship to CSF

## 4. Discussion

The primary motivation of the current study was to investigate whether two distinct pathological mechanisms of OSA, namely hypoxic burden and sleep fragmentation, are differentially associated with fluid dynamics that support glymphatic activity. Given that hypoxic injury to cerebral vasculature may promote vessel stiffening, we hypothesized that greater hypoxic burden would be associated with disrupted coordination between brain pulsations and CSF flow.^14,15^ Supporting this hypothesis, greater hypoxic burden was robustly associated with weaker gBOLD-CSF coupling, independent of non-hypoxic sleep fragmentation, hypertension, and demographic characteristics. Interestingly, neither RERAs, which represent non-hypoxic sleep fragmentation, nor AHI3%A, which captures sleep fragmentation regardless of hypoxemia, were associated with coordination between brain pulsations and CSF flow, despite the potential impact of sleep fragmentation on norepinephrine tone, which modulates vasomotion.^14,24,40^ Together, these findings suggest that hypoxic burden may be the primary driver of deficits in fluid dynamics that support glymphatic activity in OSA. Prior research has shown that coordination between vasomotion and CSF flow is a critical driver of CSF-ISF exchange and solute transport within the glymphatic system.^14,15^ Accordingly, brain pulsations and vasomotion show tight anti-phase coordination with CSF movement in normal brains.^13-15^ In comparison, weaker anti-phase coordination between brain pulsations and CSF flow has been linked to ADRD pathology.^36,41,42^ Our study extends this literature by showing that anti-phase coordination between brain pulsations and CSF flow is weakened in OSA as well, and further suggests that hypoxic burden may play a distinct role in impairing fluid dynamics that support glymphatic activity, independent of sleep fragmentation. Weakened coordination between brain pulsations and CSF flow may represent a pathway toward elevated ADRD risk in this population. To the best of our knowledge, this is the first study to disentangle the contributions of hypoxic burden and non-hypoxic sleep fragmentation to coordination between brain pulsations and CSF flow.

In addition to coordination between brain pulsations and CSF flow, brain pulsation strength may play a distinct role in glymphatic activity by inducing convective CSF flow and modulating perivascular space size.^13,15^ In healthy brains, greater brain pulsation strength has been associated with greater CSF movement and dynamic changes in perivascular space size. Contrary to our expectations, however, greater global brain pulsation strength, measured as gBOLD-amp, was associated with greater hypoxic burden, quantified using T90. Regional analyses using rBOLD-amp showed that brain pulsation strength was significantly associated with T90 across most temporal lobe regions and multiple frontal and parietal regions, but not within the occipital cortex. Effect sizes were particularly large in temporal regions, and no region showed a negative association with hypoxic burden. In contrast, sleep fragmentation, measured using RERAs, was not associated with brain pulsation strength either globally or in any cortical region. Taken together, these findings suggest that hypoxic burden in OSA, but not sleep fragmentation, predicts elevated brain pulsation strength, particularly in temporal regions. Although the mechanisms underlying this association remain unclear, the large effect sizes observed in the temporal lobe, a region with high metabolic demand that can lead to increased oxygen demand-supply mismatch in OSA, may suggest a pathological rather than adaptive process.^43,44^

Given that T90-linked elevations in brain pulsation strength were unexpected, we next examined whether these regional elevations were accompanied by stronger coordination with CSF inflow within the same regions. Prior research in healthy brains has shown that greater vasomotion is associated with stronger regional coordination with CSF movement.^14^ Interestingly, elevated brain pulsation strength did not appear to translate into improved coordination with CSF movement in “T90-susceptible regions”, which included most temporal lobe regions and multiple frontal and parietal regions showing T90-linked elevations in rBOLD-amp. In these regions, greater rBOLD-amp was not associated with stronger region-specific rBOLD-CSF coupling. This pattern contrasts with the strong association observed between rBOLD-amp and rBOLD-CSF coupling in “non-T90-susceptible regions”, consistent with previous studies showing that greater brain pulsations are tightly coupled in time with larger-magnitude CSF movement to maintain constant intracranial volume, as posited by the Monro-Kellie doctrine.^12,13,15^ Collectively, these findings suggest that regions more susceptible to hypoxic burden may exhibit greater discoordination between brain pulsations and CSF flow. Hypoxic injury to the vasculature is known to promote vascular stiffening,^20^ which may impair the ability of vascular pulsations to generate appropriately timed CSF displacement, thereby contributing to the observed dissociation between elevated region-specific brain pulsation strength and region-specific BOLD-CSF coupling. Future research is needed to determine whether elevations in brain pulsation strength linked to T90 represent pathological responses to hypoxic injury, potentially indicating impaired maintenance of stable vascular tone or increased vasodilation to preserve oxygen delivery.^21,44-46^

This study has multiple strengths. First, it distinguishes the association of hypoxic burden with fluid dynamics that support glymphatic activity from that with non-hypoxic sleep fragmentation using objective, validated sleep measures. Second, the sample included racially diverse individuals, improving generalizability to the population experiencing OSA. Third, MRI was conducted at 7T, providing improved resolution that can help enhance signal quantification, registration, and gray matter segmentation. Fourth, the OSA group consists of individuals newly diagnosed with OSA who has yet to begin treatment, precluding potential confounding by OSA treatment history.

However, this study is not without limitations. First, brain pulsations and their coordination with CSF flow were not assessed during sleep, when glymphatic activity is thought to be most active.^3,14,47^ The wakefulness-based acquisition of resting-state fMRI may explain why RERAs were not associated with deficits in brain pulsations or their coordination with CSF flow, as RERAs are likely to influence brain pulsations primarily by disrupting norepinephrine tone, an effect that may be pronounced during sleep. Nonetheless, growing evidence suggests that the glymphatic system is also active during wakefulness,^48^ and our findings show that greater hypoxic burden is associated with abnormalities in brain pulsation and CSF flow dynamics that support glymphatic function during wakefulness. Second, T90, RERAs, and AHI3%a were assessed using two different sleep study modalities, i.e., WatchPAT and in-lab polysomnography, which may have introduced measurement variability. Nonetheless, all multivariable models adjusted for sleep study modality. In addition, prior studies, together with our supplementary data, show that WatchPAT demonstrates strong agreement with in-lab polysomnography for the OSA characteristics examined in the current study.^25-27,49^ Third, our study included a modest sample size of 36 individuals. Fourth, severe hypoxemic respiratory events may co-occur with arousals. Although we additionally examined the association between gBOLD-CSF coupling and AHI3%a, which captures respiratory event-related arousals regardless of oxygen desaturation, AHI3%a is not a pure arousal metric. To address these limitations, future studies should examine larger samples assessed using a single sleep study modality, additionally evaluate arousal index as a direct measure of sleep fragmentation that includes both hypoxic and non-hypoxic arousals, and measure brain pulsations and CSF coordination during both sleep and wakefulness.

### 4.1. Conclusions

The current study shows that OSA is associated with deficits in brain pulsation and CSF flow dynamics that support glymphatic activity, with hypoxic burden emerging as the primary driver. Specifically, hypoxic burden was robustly associated with weaker coordination between global brain pulsations and CSF flow, independent of non-hypoxic sleep fragmentation, hypertension, and demographic characteristics. Greater hypoxic burden was also associated with elevated brain pulsation strength across most temporal lobe regions and multiple frontal and parietal regions. Within these regions showing T90-linked elevation in brain pulsation strength, greater brain pulsation did not translate into stronger coordination with CSF flow, deviating from the expected anti-phase coupling between blood and CSF movement under the doctrine of constant intracranial volume. The observed discoordination between brain pulsations and CSF flow may indicate a deficit in glymphatic activity, particularly in the temporal lobe, a region with high metabolic demand and vulnerability to oxygen supply-demand mismatch in OSA. In contrast, neither non-hypoxic RERAs nor AHI3%a, which captures respiratory event-related arousals regardless of hypoxemia, was associated with brain pulsation-CSF dynamics. In sum, hypoxic burden may represent a distinct pathway through which OSA contributes to impaired glymphatic activity and, in turn, ADRD risk.

## Acknowledgement of NIH funding

This work was supported by the National Institutes of Health under award number R01 AG066870 and R01 AG080609. This manuscript is the result of funding in whole or in part by the National Institutes of Health (NIH). It is subject to the NIH Public Access Policy. Through acceptance of this federal funding, NIH has been given a right to make this manuscript publicly available in PubMed Central upon the Official Date of Publication, as defined by NIH.

## Notes

**FUNDING.**This work was supported by the National Institute of Health (R01 AG066870, R01 AG080609) and the Alzheimer’s Association (AARFD-24-1306796).

### Competing Interest Statement

The authors have declared no competing interest.

### Summary of Updates

Additional exploratory analysis and Results section updated.

