## Supplementary Files for "Greater hypoxic burden predicts weaker coordination between brain pulsation and CSF flow on 7T MRI independent of non-hypoxic arousals: Implications for glymphatic activity"

**Supplementary Figure 1.** Participant flow diagram

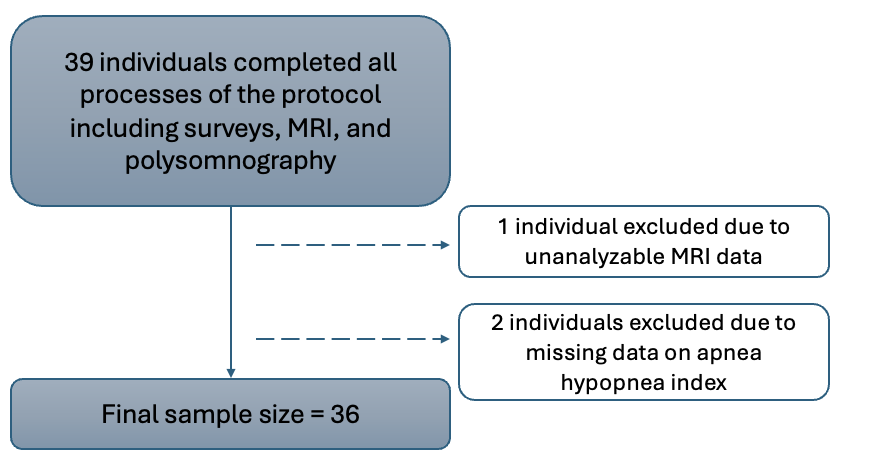

**Supplementary Table 1.** Agreement between WatchPAT and in-lab polysomnography

| **Characteristics (WatchPAT vs. in-lab polysomnography)** | **ICC** | **F^1^** | ***p*** |
| --- | --- | --- | --- |
| pRDI vs. AHI3%a | 0.76 | 4.21 | <0.001 |
| pAHI3 vs. AHI3% | 0.76 | 4.08 | <0.001 |

^1^ F-test for statistical difference of ICC from 0

*Acronyms.* AHI3%= the sum of all apneas and hypopneas with amplitude of breathing by 30% or more for ≥ 10 seconds terminating in ≥ 3% decline in blood oxygen saturation or arousals as assessed by in-lab polysomnography; AHI3%a=AHI3% +event-related arousals as assessed by in-lab polysomnography; pAHI3=AHI3=WatchPAT-derived AHI3%; pRDI=WatchPAT-derived respiratory disturbance index

**Supplementary Table 2.** Association between rBOLD-amp and T90

| **Region of interest** | **r** | ***P***^1^ |
| --- | --- | --- |
| ctx_lh_bankssts | 0.32 | 0.06 |
| ctx_lh_caudalanteriorcingulate | 0.35 | 0.04 |
| ctx_lh_caudalmiddlefrontal | 0.24 | 0.16 |
| ctx_lh_cuneus | 0.25 | 0.14 |
| ctx_lh_entorhinal | 0.08 | 0.64 |
| ctx_lh_fusiform | 0.33 | 0.05 |
| ctx_lh_inferiorparietal | 0.25 | 0.15 |
| ctx_lh_inferiortemporal | 0.18 | 0.30 |
| ctx_lh_isthmuscingulate | 0.29 | 0.09 |
| ctx_lh_lateraloccipital | 0.25 | 0.14 |
| ctx_lh_lateralorbitofrontal | 0.39 | 0.02 |
| ctx_lh_lingual | 0.30 | 0.07 |
| ctx_lh_medialorbitofrontal | 0.34 | 0.04 |
| ctx_lh_middletemporal | 0.35 | 0.04 |
| ctx_lh_parahippocampal | 0.23 | 0.18 |
| ctx_lh_paracentral | 0.35 | 0.04 |
| ctx_lh_parsopercularis | 0.32 | 0.06 |
| ctx_lh_parsorbitalis | 0.43 | 0.01 |
| ctx_lh_parstriangularis | 0.25 | 0.14 |
| ctx_lh_pericalcarine | 0.23 | 0.19 |
| ctx_lh_postcentral | 0.39 | 0.02 |
| ctx_lh_posteriorcingulate | 0.29 | 0.09 |
| ctx_lh_precentral | 0.32 | 0.06 |
| ctx_lh_precuneus | 0.24 | 0.16 |
| ctx_lh_rostralanteriorcingulate | 0.39 | 0.02 |
| ctx_lh_rostralmiddlefrontal | 0.32 | 0.06 |
| ctx_lh_superiorfrontal | 0.38 | 0.02 |
| ctx_lh_superiorparietal | 0.19 | 0.27 |
| ctx_lh_superiortemporal | 0.44 | 0.01 |
| ctx_lh_supramarginal | 0.26 | 0.13 |
| ctx_lh_frontalpole | 0.38 | 0.03 |
| ctx_lh_temporalpole | 0.40 | 0.01 |
| ctx_lh_transversetemporal | 0.38 | 0.02 |
| ctx_lh_insula | 0.42 | 0.01 |
| ctx_rh_bankssts | 0.31 | 0.06 |
| ctx_rh_caudalanteriorcingulate | 0.27 | 0.11 |
| ctx_rh_caudalmiddlefrontal | 0.15 | 0.39 |
| ctx_rh_cuneus | 0.25 | 0.14 |
| ctx_rh_entorhinal | 0.22 | 0.19 |
| ctx_rh_fusiform | 0.34 | 0.04 |
| ctx_rh_inferiorparietal | 0.21 | 0.21 |
| ctx_rh_inferiortemporal | 0.44 | 0.01 |
| ctx_rh_isthmuscingulate | 0.20 | 0.25 |
| ctx_rh_lateraloccipital | 0.27 | 0.12 |
| ctx_rh_lateralorbitofrontal | 0.37 | 0.03 |
| ctx_rh_lingual | 0.32 | 0.06 |
| ctx_rh_medialorbitofrontal | 0.25 | 0.13 |
| ctx_rh_middletemporal | 0.39 | 0.02 |
| ctx_rh_parahippocampal | 0.38 | 0.02 |
| ctx_rh_paracentral | 0.22 | 0.19 |
| ctx_rh_parsopercularis | 0.31 | 0.07 |
| ctx_rh_parsorbitalis | 0.33 | 0.05 |
| ctx_rh_parstriangularis | 0.28 | 0.10 |
| ctx_rh_pericalcarine | 0.17 | 0.33 |
| ctx_rh_postcentral | 0.31 | 0.06 |
| ctx_rh_posteriorcingulate | 0.16 | 0.35 |
| ctx_rh_precentral | 0.29 | 0.09 |
| ctx_rh_precuneus | 0.17 | 0.33 |
| ctx_rh_rostralanteriorcingulate | 0.36 | 0.03 |
| ctx_rh_rostralmiddlefrontal | 0.30 | 0.07 |
| ctx_rh_superiorfrontal | 0.31 | 0.07 |
| ctx_rh_superiorparietal | 0.22 | 0.19 |
| ctx_rh_superiortemporal | 0.43 | 0.01 |
| ctx_rh_supramarginal | 0.28 | 0.09 |
| ctx_rh_frontalpole | 0.05 | 0.77 |
| ctx_rh_temporalpole | 0.29 | 0.09 |
| ctx_rh_transversetemporal | 0.34 | 0.04 |
| ctx_rh_insula | 0.41 | 0.01 |

^1^*p*-values were derived based on permutation testing.

Acronyms. T90=time spent under 90% oxygen saturation

**Supplementary Table 3.** Association between rBOLD-amp and RERAs

| **Region of interest** | **r** | ***P***^1^ |
| --- | --- | --- |
| ctx_lh_bankssts | -0.15 | 0.39 |
| ctx_lh_caudalanteriorcingulate | -0.10 | 0.57 |
| ctx_lh_caudalmiddlefrontal | 0.06 | 0.73 |
| ctx_lh_cuneus | -0.10 | 0.59 |
| ctx_lh_entorhinal | -0.17 | 0.34 |
| ctx_lh_fusiform | -0.14 | 0.42 |
| ctx_lh_inferiorparietal | -0.23 | 0.19 |
| ctx_lh_inferiortemporal | -0.05 | 0.78 |
| ctx_lh_isthmuscingulate | -0.02 | 0.90 |
| ctx_lh_lateraloccipital | -0.03 | 0.88 |
| ctx_lh_lateralorbitofrontal | 0.02 | 0.92 |
| ctx_lh_lingual | -0.01 | 0.97 |
| ctx_lh_medialorbitofrontal | -0.18 | 0.31 |
| ctx_lh_middletemporal | 0.05 | 0.78 |
| ctx_lh_parahippocampal | -0.08 | 0.64 |
| ctx_lh_paracentral | -0.13 | 0.44 |
| ctx_lh_parsopercularis | -0.13 | 0.45 |
| ctx_lh_parsorbitalis | -0.18 | 0.29 |
| ctx_lh_parstriangularis | -0.01 | 0.94 |
| ctx_lh_pericalcarine | -0.20 | 0.26 |
| ctx_lh_postcentral | -0.10 | 0.57 |
| ctx_lh_posteriorcingulate | -0.20 | 0.25 |
| ctx_lh_precentral | -0.01 | 0.97 |
| ctx_lh_precuneus | -0.10 | 0.58 |
| ctx_lh_rostralanteriorcingulate | -0.15 | 0.38 |
| ctx_lh_rostralmiddlefrontal | -0.10 | 0.55 |
| ctx_lh_superiorfrontal | -0.01 | 0.95 |
| ctx_lh_superiorparietal | -0.21 | 0.21 |
| ctx_lh_superiortemporal | -0.23 | 0.19 |
| ctx_lh_supramarginal | 0.06 | 0.74 |
| ctx_lh_frontalpole | -0.01 | 0.96 |
| ctx_lh_temporalpole | -0.30 | 0.08 |
| ctx_lh_transversetemporal | -0.15 | 0.38 |
| ctx_lh_insula | 0.03 | 0.86 |
| ctx_rh_bankssts | -0.08 | 0.65 |
| ctx_rh_caudalanteriorcingulate | -0.16 | 0.37 |
| ctx_rh_caudalmiddlefrontal | -0.01 | 0.96 |
| ctx_rh_cuneus | 0.11 | 0.52 |
| ctx_rh_entorhinal | -0.03 | 0.89 |
| ctx_rh_fusiform | 0.08 | 0.63 |
| ctx_rh_inferiorparietal | 0.09 | 0.59 |
| ctx_rh_inferiortemporal | -0.01 | 0.98 |
| ctx_rh_isthmuscingulate | 0.03 | 0.87 |
| ctx_rh_lateraloccipital | -0.06 | 0.74 |
| ctx_rh_lateralorbitofrontal | 0.03 | 0.88 |
| ctx_rh_lingual | -0.05 | 0.76 |
| ctx_rh_medialorbitofrontal | -0.04 | 0.83 |
| ctx_rh_middletemporal | 0.15 | 0.39 |
| ctx_rh_parahippocampal | 0.02 | 0.93 |
| ctx_rh_paracentral | -0.18 | 0.31 |
| ctx_rh_parsopercularis | -0.19 | 0.27 |
| ctx_rh_parsorbitalis | -0.23 | 0.18 |
| ctx_rh_parstriangularis | 0.02 | 0.91 |
| ctx_rh_pericalcarine | -0.08 | 0.66 |
| ctx_rh_postcentral | -0.04 | 0.84 |
| ctx_rh_posteriorcingulate | -0.10 | 0.56 |
| ctx_rh_precentral | 0.03 | 0.88 |
| ctx_rh_precuneus | -0.09 | 0.62 |
| ctx_rh_rostralanteriorcingulate | -0.24 | 0.17 |
| ctx_rh_rostralmiddlefrontal | -0.15 | 0.40 |
| ctx_rh_superiorfrontal | 0.03 | 0.87 |
| ctx_rh_superiorparietal | -0.05 | 0.80 |
| ctx_rh_superiortemporal | -0.003 | 0.99 |
| ctx_rh_supramarginal | -0.16 | 0.35 |
| ctx_rh_frontalpole | <0.01 | 0.98 |
| ctx_rh_temporalpole | -0.07 | 0.72 |
| ctx_rh_transversetemporal | -0.04 | 0.81 |
| ctx_rh_insula | -0.15 | 0.39 |

^1^*p*-values were derived based on permutation testing.

Acronyms. RERA=respiratory effort-related arousals
